# Genome-wide Estrogen Receptor-*α* activation is sustained, not cyclical

**DOI:** 10.1101/398925

**Authors:** Andrew N Holding, Amy E Cullen, Florian Markowetz

## Abstract

Estrogen Receptor-*α* (ER) is the key driver of 75% of all breast cancers. Upon stimulation by its ligand estra-2-diol, ER forms a transcriptionally active complex binding chromatin. Previous studies have reported that ER binding follows a cyclical binding pattern with a periodicity of 90 minutes. However, these studies have been limited to individual ER target genes and most were done without replicates. Thus, the robustness and generality of ER cycling are not well understood.

Here we present a comprehensive genome-wide analysis of the time dependence of ER binding affinity up to 90 minutes after activation, based on 6 replicates at 10 time points using our previously reported method for precise quantification of binding, Parallel-Factor ChIP-seq (pfChIP-seq). In contrast to previously described cyclical binding, our approach identifies a unidirectional sustained increase in ER binding affinity, as well as a class of estra-2-diol independent binding sites. Our results are corroborated by a quantitative re-analysis of data from multiple independent studies.

Our new model reconciles the results of multiple conflicting studies into the activation of ER at the TFF1 promoter. We provide a detailed understanding of ER’s response to estra-2-diol in the context of the receptor’s fundamental role as both the main driver and therapeutic target of breast cancer.

## 1 Introduction

The study of the Estrogen Receptor-*α* (ER) has played a fundamental role in both our understanding of transcription factors and cancer biology. The ER is one of a family of transcription factors called nuclear receptors. Nuclear receptors are intra-cellular and, on activation by their ligand, typically undergo dimerisation and bind to specific DNA motif (for ER: Estrogen Response Elements; EREs). On the chromatin, the nuclear receptor recruits a series of cofactors and promotes the basal transcription mechanism at either nearby promoters or through chromatin loops from distal enhancers. Because of the minimal nature of these systems relative to other signaling pathways, nuclear receptors have become a model system for transcription factor analysis. Simultaneously, the role of nuclear receptors as drivers in a range of hormone dependent cancers has led to focused studies in the context of the disease.

Previously, it was reported that the ER and key cofactors followed a cyclical pattern in breast cancer cell lines with maximal binding at 45 minutes after stimulation with estra-2-diol [1, 2]. Similar results were also reported for the AR after activation with DHT [3] and several follow-up studies exist looking at single genomic *loci* [4, 5, 6, 7]. However, subsequent genome-wide studies have provided little further detail on the specific nature of the proposed kinetics of ER binding being either limited in the number of replicates or lacking temporal resolution [8, 9, 10, 11]. In our own network analysis [12], we focused on 0, 45 and 90 minutes and found no significant reduction in ER signal at 90 minutes. In the same study, quantitative proteomic analysis of ER interactions at the same time intervals by qPLEX-RIME [13] shows no significant difference in terms of ER interactions at 45 and 90 minutes. These conflicting results have so far not been resolved.

Routinely used assays to measure protein binding to chromatin are based on Chromatin Immunoprecipitation (ChIP). A major challenge to monitoring ER activation through ChIP is the normalization of the ChIP signal — either genome-wide with next generation sequencing or at individual loci by qPCR — as the standard protocols do not control for a significant number of confounding factors including the efficiency of the immunoprecipitation step. In the case the of the two original studies [1, 2], the data only provided limited controls in this regard. An alternative method that has been applied to normalize ChIP-seq data is to use the maximal read count obtained at each individual site across each time point [11]; however, this method is at the expense of monitoring the magnitude of ER binding and gives equal weight to low read count peaks and more robust data from stronger binding sites.

In the context of these challenges, we applied two strategies to robustly and accurately monitor the process of nuclear receptor binding to chromatin on activation. The first strategy was to increase the number of replicates. We generated sample data for six independent isogenic experiments to enable better characterization of the variance within the data. This strategy provided an unprecedented level of information regarding ER activation with twice the level of replication used in previous ChIP-qPCR studies [2] and a significant improvement on previous single replicate genome-wide studies. The second strategy was to use our recently developed method for precise quantification of binding, Parallel-Factor ChIP (pfChIP) [14], which uses an internal control for quantitative differential ChIP-seq [3]. Combined, these two strategies enabled us to undertake the most comprehensive and precise analysis of ER activation to date.

## 2 Results

### Measurement of Genome Copy-Number Discordance

We measured ER-binding in MCF7 cells, a widely used model system for ER biology. To maximize the reproducibility of our results, MCF7 cells were grown from ATCC stocks, keeping passaging to a minimum, and the cell line origin was confirmed by STR genotyping. Additionally, to ensure the MCF7 cell line did not show significant genetic drift during culturing within our laboratory, we applied Cell-Strainer [15] to the input data from our ChIP-seq experiments. The fraction of genome with copy-number discordance was estimated at 0.2787, within the range of 0 to 0.3 as published by CellStrainer’s developers to ensure similar therapeutic response.

### Visualization of raw data

Sequencing reads from the analysis of 60 pfChip-seq samples targeting ER and six input samples were demultiplexed and aligned to the Homo sapiens GRCh38 reference assembly. Visual inspection of the data using the Integrative Genomics Viewer (IGV) Viewer [16] confirmed enrichment at known ER binding sites (exemplified by TFF1 in Figure S1) and the presence of previously reported CTCF control peaks [14]. From visual inspection, pfChIP-seq samples qualitatively showed minimal ER binding at 0 minutes while CTCF binding was constant at all time points.

### Parallel-Factor Normalization

Peak count data from CTCF binding sites were used to normalize between conditions, with >70 000 binding sites discovered across all samples and >50 000 CTCF binding sites found in over 50% of samples. Analysis after normalization of the raw data showed similar levels of variability in terms of signal (Figure S2) as we saw when developing the pfChIP method [14]. The resultant normalized binding matrix of ER binding was used for all downstream analyses and is provided as Supplementary Table 1.

### ER Binding at the TFF1 Promoter

Normalized count data for the TFF1 promoter showed that on activation with estra-2-diol the ER rapidly (in less than 10 minutes) binds the TFF1 promoter. Binding after this time point shows no significant changes. Analysis of the data by individual replicates (Figure S3) did not demonstrate evidence of oscillatory binding in individual replicates either with a period of 90 minutes period or an alternative frequency.

Comparison of the variance in the ER binding after induction shows that there is significantly more variance (F-test, time points >= 10 minutes, p-value < 1 × 10^−10^) in the ER binding data than in CTCF binding between replicates. As the variance of CTCF binding in pfChIP-seq is a good estimator of the technical variance, the most likely source of increased variance in ER binding is therefore biological. These findings were validated through analysis of the RARA promoter and proximal CTCF peaks (Figure S4), which gave consistent results to those seen at the TFF1 promoter.

### Locus Specific Variation in Maximal ER Binding Affinity

Previously, ER binding sites were shown to reach maximum occupancy at different time points depending on genomic location, revealing a P300 squelching mechanism at early time points [11]. Therefore, to provide a partial validation of this study, we applied the same principles of their analysis to our data, i.e. normalizing in the time-space setting maximum occupancy to 1. Consistent with the previous study, the two time points with the largest numbers of sites reaching maximal occupancy in both data sets were at 10 and 40 minutes (Figure 2A). As the remaining time points were unique to the individual data sets, these could not be directly compared.

**Figure 1:**
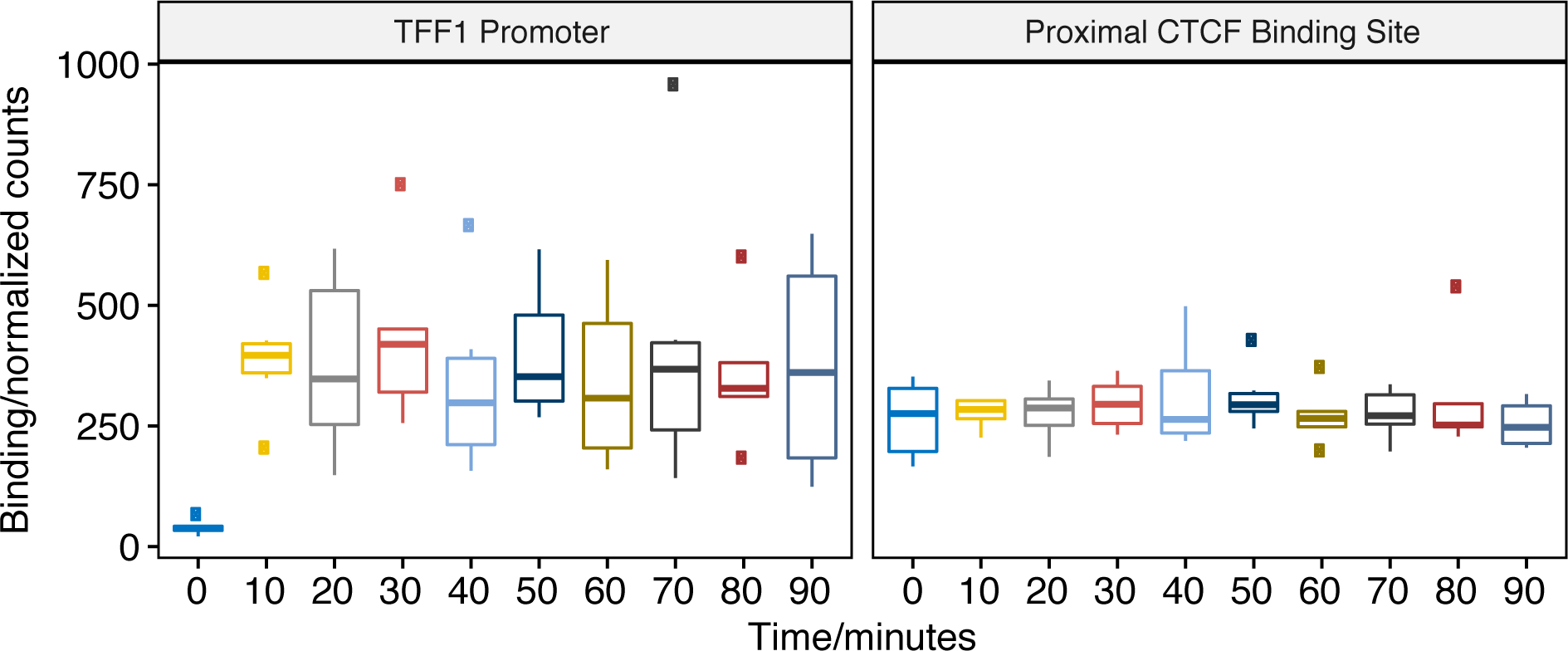
pfChIP-seq signal at the TFF1 promoter and proximal CTCF binding site. Binding of ER at the TFF1 promoter has been the classical focus of study before genome-wide technology and the predicted site for oscillations in ER binding. ER binding is minimal at 0 minutes; however, by 10 minutes, the ER has rapidly and robustly bound to give a sustained signal at the TFF1 promoter. In contrast, the closest CTCF binding site demonstrates a constant, estra-2-diol-independent, signal with significantly less variance.

**Figure 2:**
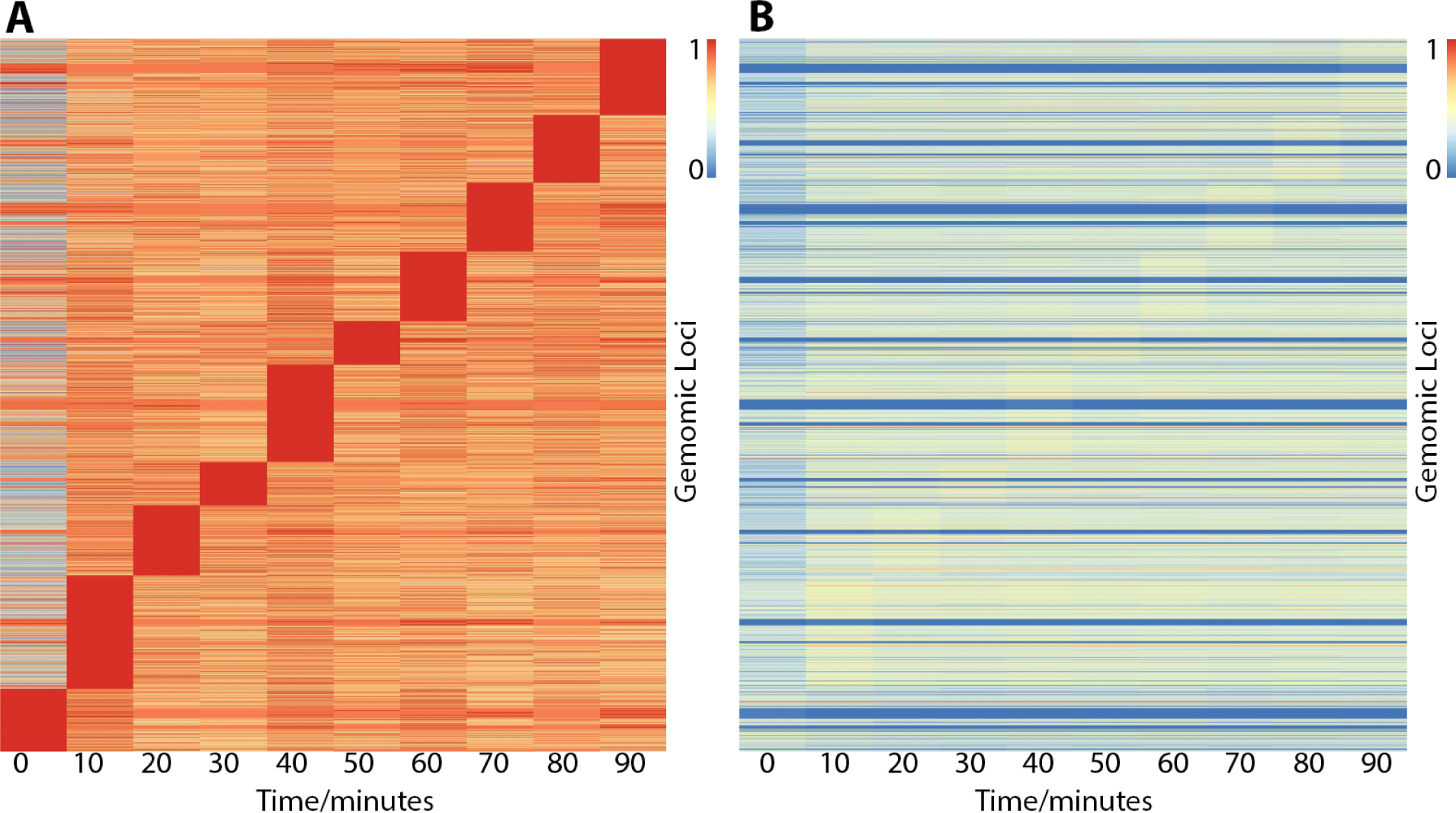
Heatmaps showing ER binding affinity from 0 to 90 minutes after stimulation with estra-2-diol normalized in two different ways. Row order is the same in both plots. (A) Normalized by row to time point with maximal binding. Data suggests that genomic loci may influence the time point maximal binding; however, normalizing to CTCF control peaks (B) demonstrates the effect is potentially overemphasized by normalization choice and that binding affinity is the biggest variable. In contrast, both plots (A and B) show minimal ER binding affinity is found at 0 minutes, consistent with the literature response of MCF7 cells to treatment of estra-2-diol.

Furthermore, pfChIP-seq allowed us to improve on the previous study by directly normalizing the data to the internal control. The resultant binding matrix provided quantification of the absolute binding affinity at each time point (Figure 2B).

Comparison of Figures 2A and 2B demonstrates the effects of different data normalization strategies. The relative normalization to maximum binding emphasizes binding maxima (red blocks in Figure 2A) while the absolute normalization to an internal control shows that these maxima are very shallow, barely visible in Figure 2B, and other features dominate the data. A few genes show very high levels of ER binding (visible as thin red lines in Figure 2B), while most genes show intermediate levels and some very low levels (blue lines). These different levels of ER binding are preserved over time, with only time point 0 showing very low levels for all genes.

### Visualizing Temporal ER Binding Affinity

To elucidate potential different temporal responses to ER activation by estra-2-diol, we applied t-SNE [17], a widely used method for dimensionality reduction and data visualization (Figure 3). Each dot in the plot represents a binding site over time, i.e. one row in the binding matrix shown in Figure 2B. We colored each dot by the false discovery rate (FDR; [18]) for the change in ER affinity between 0 to 10 minutes. This analysis revealed two major trajectories of binding sites in the data, one dominated by low FDR (orange) and one by high FDR (blue). Both trajectories saw an increasing affinity in the direction of the white arrow. This pattern was stable for a wide range of perplexity, the main t-SNE parameter (Supplementary Figure S5).

**Figure 3:**
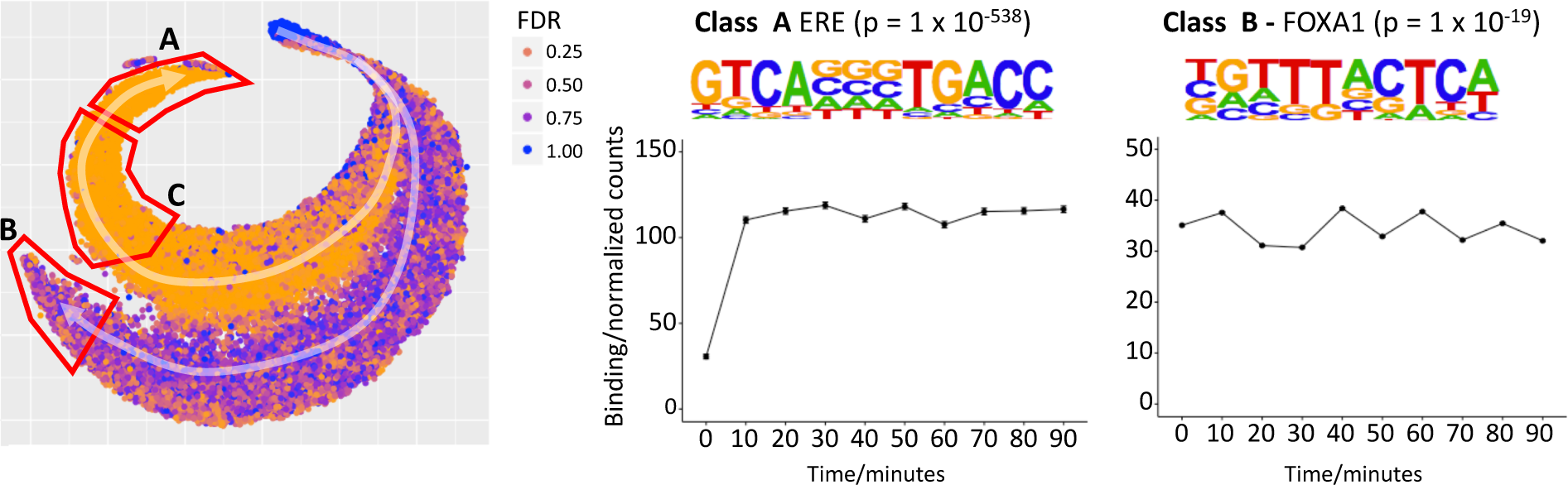
t-SNE plot to explore temporal patterns in ER binding affinity. Two trajectories, A and B, are highlighted with white arrows and starting a single cluster of peaks. Points are colored by FDR value computed by Brundle/DiffBind for the 0 vs 10 minutes contrast. Trajectory A demonstrates increasing ER affinity in response to estra-2-diol at 10 minutes. Trajectory B shows increasing affinity for all times points, i.e. estra-2-diol independent binding, but the maximum signal is of a lower intensity than that of Trajectory A. *De novo* motif analysis for Class A (the peaks found at the end of trajectory A) gave strongest enrichment for the ERE (p = 1×10^−538^). The same analysis of Class C provided a partial ERE (not shown), consistent with ER affinity being a function of how conserved the ER binding site is with respect to ideal ERE. Analysis of Class B gave FOXA1 as the most significantly enriched motif (p = 1 × 10^−19^).

We named the estra-2-diol responsive trajectory A, and the estra-2-diol independent trajectory B. The set of genomic sites found at the end of each trajectory were named Class A and B respectively. Motif analysis of Class A peaks demonstrated significant enrichment for the full estrogen response element (ERE, [19]), while Class B gave enrichment for the *FOXA1* binding site. Analysis of Class C (i.e. weaker responding genes on trajectory A) gave a partial ERE match, suggesting a greater divergence from the ERE motif and consistent with the lower levels of ER affinity found on ER activation at these sites [20].

Average binding profiles were computed for both Class A and Class B. Class A showed minimal binding at 0 minutes followed by a robust response before 10 minutes, the binding affinity then remained similar for the remaining time points. In contrast, Class B displayed estra-2-diol independent binding at 0 minutes and average ER binding affinity saw no significant changes between time points. Class C gave a similar profile to Class A (not shown), but with reduced amplitude. The average amplitude of the binding from 10–90 minutes displayed a greater ER affinity for Class A then Class B.

Genomic regions enrichment of annotations tool (GREAT) analysis [21] of Class B binding sites (Supplementary Table 2) identified the enrichment of six amplicons previously identified from the analysis of 191 breast tumor samples, q = 5.6 × 10^−41^ to q = 3.3×10^−8^, [22] and a set of genes upregulated in luminal-like breast cancer cell lines compared to the mesenchymal-like cell lines, q = 1.9 × 10^−13^, [23]).

### Analysis of Class A ER Binding Sites

Class A binding sites showed the strongest response to estra-2-diol, the greatest enrichment of the estrogen response element and contained the classical ER binding site at TFF1. We therefore focused further analysis on these peaks to minimize confounding factors. A t-SNE plot of only Class A sites (Figure 4) did not provide distinct clustering of points. Partial separation was seen on the basis of time point of maximal binding (left to right) and amplitude (approximately top to bottom).

**Figure 4:**
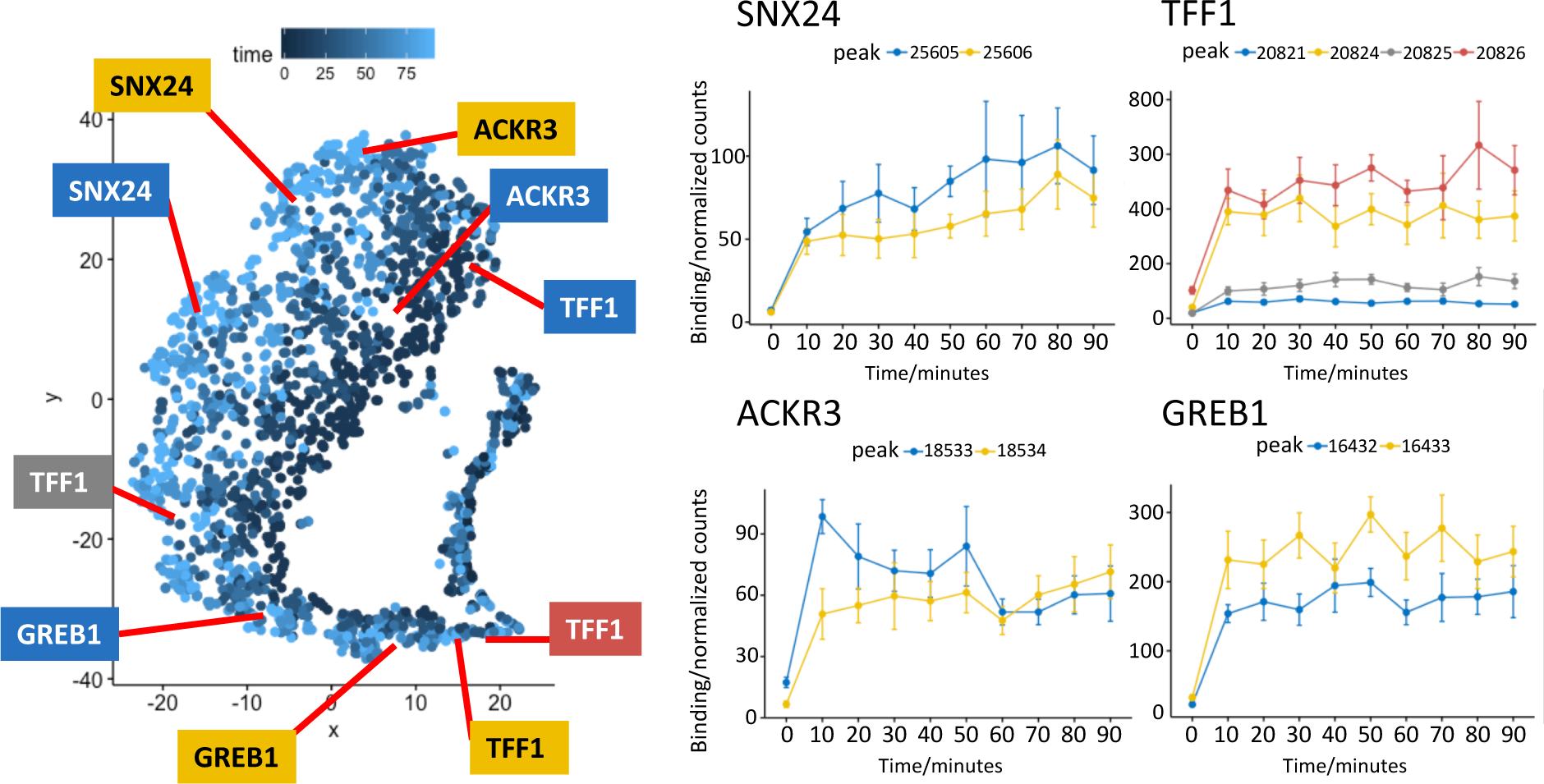
t-SNE plot of Class A from Figure 3, points colored by time of maximum value. Profiles for binding sites near the transcription start site of two well studied ER target genes, *TFF1* and *GREB1*, gave a robust sustained response to estra-2-diol. Binding sites near *SNX24* and *ACKR3* TSS are shown to examples of ER binding affinity profiles that indicate potential early or late maximal binding. Peak coordinates are provided in Supplementary Table 1.

As the class profiles may average out site-specific oscillatory kinetics, we undertook analysis of individual ER binding sites. Peaks were annotated on the basis of the nearest Transcription Start Sites (TSS) and profiles for key ER target genes *TFF1* and *GREB1* were generated. As previously seen in Figure S1, ER binding at TFF1 was stable after induction. The same response was seen at the TFF1 enhancer (dark red). Analysis of ER binding proximal to *GREB1* again showed a robust and unidirectional response to estra-2-diol.

Profiles of ER binding that showed either early or late maximal ER affinity were individually investigated. Binding near the TSS of *SNX24* and *ACKR3* are provided as representative examples.

### Quantitative re-analysis of independent studies

Given we found a robust and stable response to ER activation by estra-2-diol in contrast to the cyclical response previously described [1], we reviewed studies that have investigated ER binding at the *TFF1* promoter. Several studies either used a different promoter [24], factor [25] or estra-2-diol concentration/include *α*-amanitin [2].

By manually reviewing the first 1000 citations of [1], we identified several studies [7, 4, 6] that undertook the same analysis in the MCF7 cell line, with the same concentration of estra-2-diol, same crosslinking time scale, and at the same promoter. Since the numerical values of ER binding occupancy were not available for these studies, we read the values off the provided charts or undertook image analysis of figures (Supplementary Table 4).

Comparison of the data from all four studies gave little or no consistency in the temporal profile of ER, AIB1 and P300 binding at these sites (Figure S6). Interpretation is further hindered as these studies only report a single replicate for analysis, thereby making it impossible to quantify uncertainty in the data. Therefore, there is no consistent evidence for cycling in the studies using the same conditions as the original observation.

## 3 Discussion

By undertaking six biological replicates and incorporating an internal control with pfChIP, we have produced the most comprehensive analysis to date of ER binding over the first 90 minutes after stimulation with estra-2-diol. We found the sites at which we detected ER binding on the chromatin follows two distinct trajectories, either the rapid activation within 10 minutes followed by a stable response or ligand independent binding.

Enrichment of the FOXA1 motif in the strongest ligand-independent/Class B sites supports our hypothesis: that these are as a result of ER interactions at these sites. Importantly, the *de novo* motif analysis did not find the presence of the CTCF motif, confirming that they are not an artifact of utilizing CTCF to normalize *via* the pfChIP-seq method. Analysis of the Class B binding sites with GREAT [21], Supplementary Table 2, gave enrichment for 6 out of 30 ER regulated amplicons identified in a previous study of 191 breast cancer tumor samples [22]. On the basis that no ERE was found at Class B sites and that the affinity of ER at these sites was less than at estra-2-diol response sites, we propose that these sites represent open regions of chromatin where ER can be recruited by other transcription factors in the absence of its own ligand. However, these interactions are weak, and very likely transient, as the average binding affinity for Class B sites is similar in level (a normalized read count of 30–40) to the binding before activation at Class A binding sites, but greater than Class C binding sites (≈10 normalized reads increasing to ≈40 on activation).

Ligand dependent activation of ER was seen robustly at Class A sites, but displayed no evidence of cyclical binding. We propose instead that ER activation occurs rapidly, within 10 minutes and binding shows no significant change after this point. The two examples we demonstrated — of increasing or decreasing ER binding after activation at the *SNX24* and *ACKR3* TSS (Figure 4) — should be interpreted with caution as, while downstream effects are likely to modulate ER binding, searching for individual outliers results within a large data set will generate false positives. Nonetheless, the two examples imply a secondary level of modulation does occur as previously seen, but at much lower magnitude than proposed in studies focused on ER cycling.

In light of our results and the lack of consistency of published results, we propose that the previously described cyclical response kinetics are likely an artefact of observing a highly variable process without replicates. With replicates, the cyclical effect is lost when averaging. Even if a cyclical response existed, our results indicate that it is not regulated tightly enough to be coherently visible across multiple replicates. The variance in ER binding may better be described by heterogeneity in the cell populations before induction and by current models regarding expression noise as an indicator for greater transcription responsiveness [26].

While we cannot discount that our cells could have specifically lost the ability to regulate ER binding in the manner previously described, we have minimized this possibility through the use of cells direct from ATCC, by confirming the cell line by STR genotype and applying the latest methods [15] to confirm that our cell line is genetically similar to the strains used in other labs. Nonetheless, we would welcome further replication of this study.

In summary, through the use of stringent internal controls, we have repro-ducibly shown that estra-2-diol responsive ER binding is sustained and not cyclical, with the magnitude of the binding primarily defined by the conservation of the ERE at the binding site.

## 4 Methods and Materials

### Cell Culture

MCF7 cells were obtained from ATCC and confirmed by STR genotype before culture. For each immunoprecipitation, cells from 2×15 cm dishes were used. In each 15 cm plate, 2×10^6^ were seeded and grown for 3 days in DMEM (Glibco) with 10% FBS before washing with phosphate buffered saline. Media was replaced with charcoal stripped and phenol red-free DMEM medium. Media was replaced daily for 4 days to ensure removal of estrogenic compounds. Plates were stimulated on day 5 with a final concentration of 100 nM estra-2-diol in EtOH before crosslinking at the required time. All six replicates were done on different dates and represent different passages.

### pfChIP-seq

Parallel-factor ChIP-seq was performed as previously described [14]. CTCF antibody was D31H2 Lot:3 (Cell Signaling). ER antibody was 06-965 Lot:3008172 (Millipore).

### Data Analysis

Reads were aligned using BWA [27], and ENCODE blacklist regions [28] were removed as previously described [29]. Duplicate reads were removed and peak calling was undertaken using MACS2 [30, 31]. ER and CTCF peaks were filtered according to the pfChIP-seq protocol[14], before normalization and differential binding analysis with Brundle/DiffBind [14, 32] in R. t-SNE plots were generated with Rtsne [33]. Perplexity was tested from 2 to 200 to confirm the stability of the transformation of the data into 2-dimensional space S5. Lower perplexities, 2 and 5, gave minimal structure. For perplexities tested between 30 and 200, two stable trajectories were seen in all cases. GREAT [21] was used to analyze Class B binding sites. Band intensities from previously published studies were measured with ImageJ [34].

### Data Repositories

All sequencing data is available from NCBI Gene Expression Omnibus. Data set reference waiting allocation.

## 5 Acknowledgments

Experimental Design, Sample Preparation and Data Analysis, ANH. Sample Preparation, AEC. Manuscript preparation ANH, AEC and FM. We thank the CRUK Genomics Core for undertaking the library preparation and sequencing.

---

**Figure S1:**
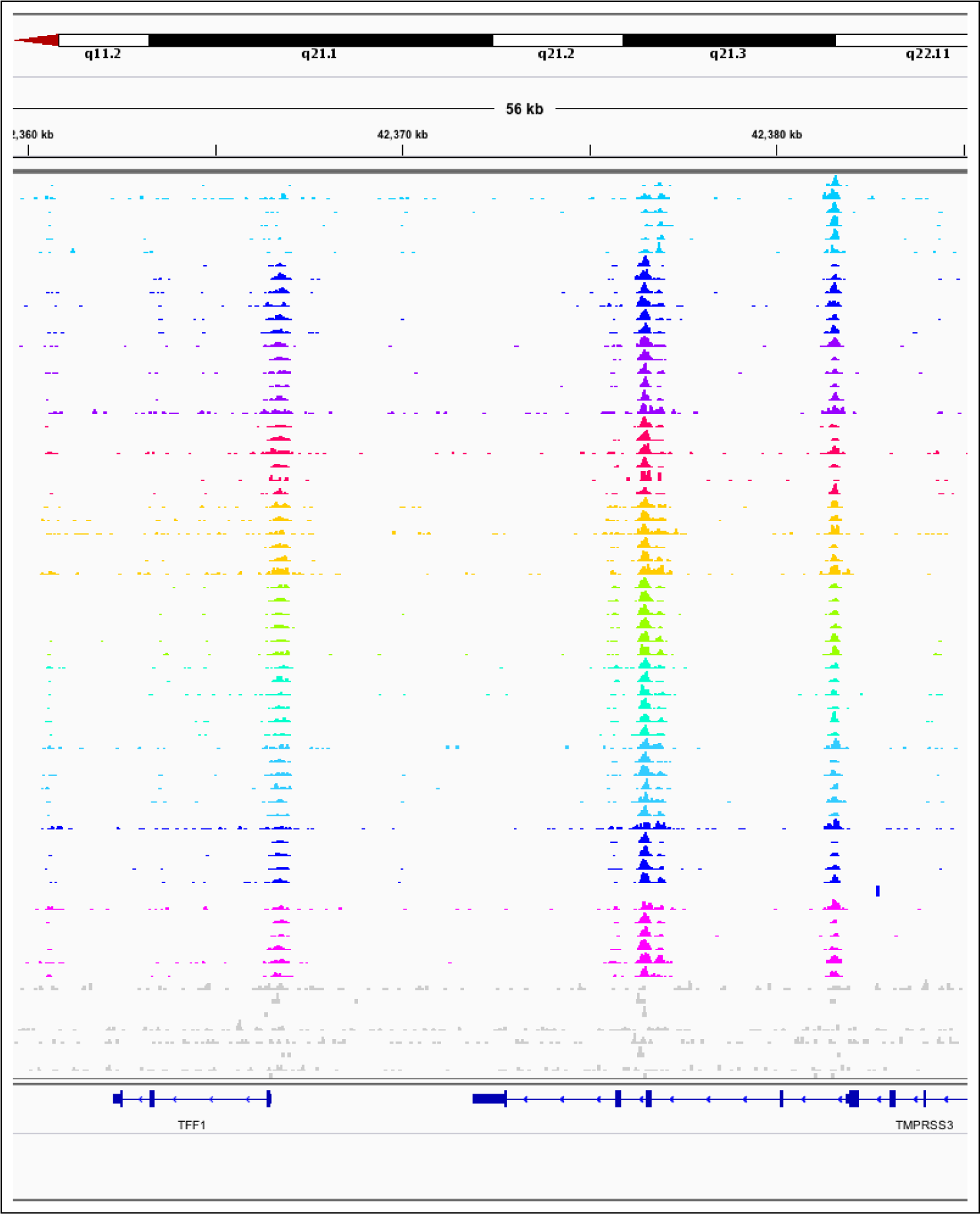
TFF1 Promoter and Enhancer. Illustrative screenshot of aligned reads from one lane of sequencing after demultiplexing. Key features are the pile-up of reads at the TFF1 promoter (left) and enhancer (middle) and the CTCF control peak (right). From top to bottom is 0 to 90 minutes in 10 minute intervals, six replicates of each. Light gray samples are input controls. Row height normalized to maximum read count in region.

**Figure S2:**
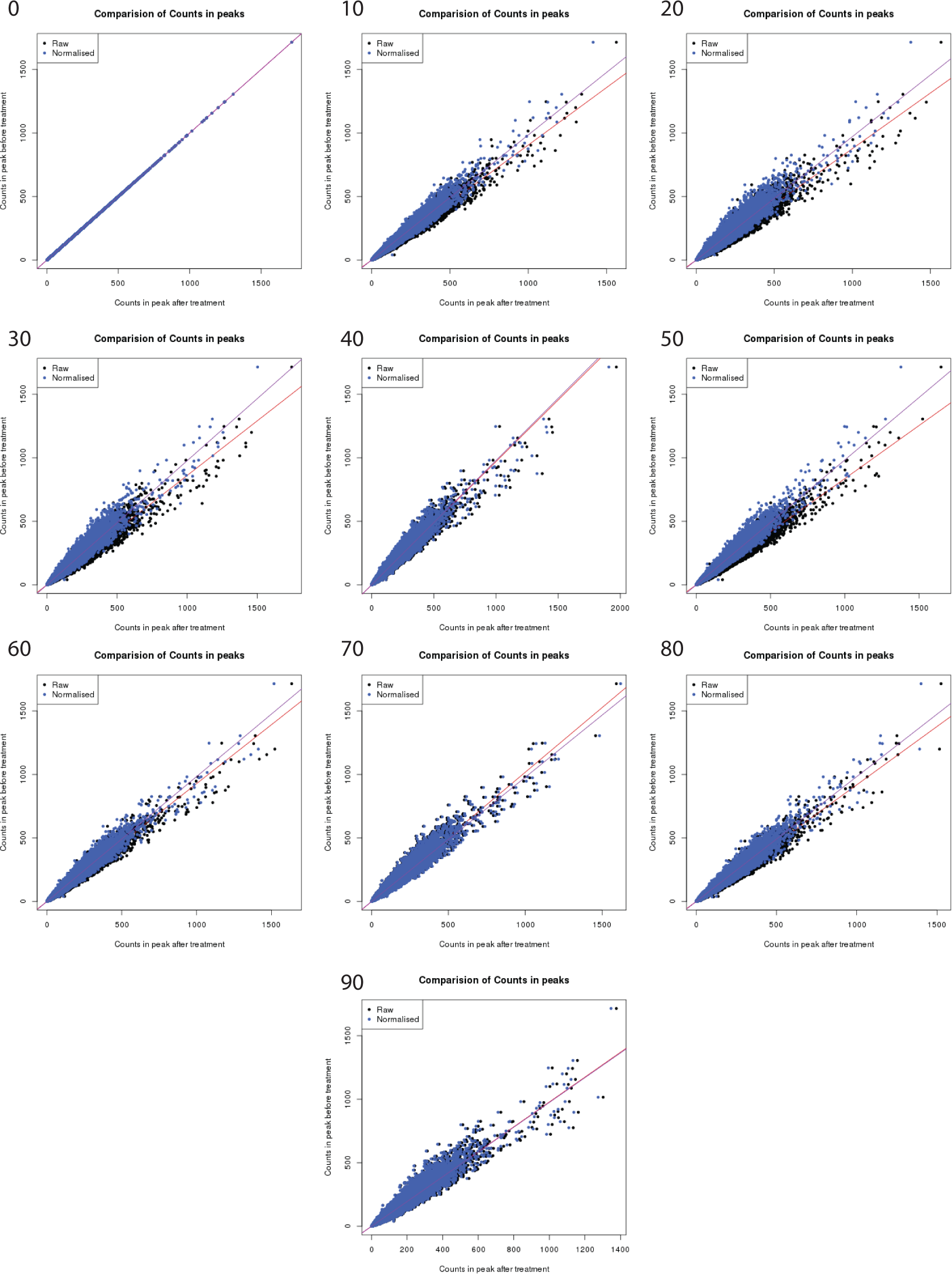
Normalization plots for each time point as generated by Brundle [14]. Each point represents a CTCF control binding site. After establishing the normalization factor for parity of CTCF binding, the ER binding was corrected using the same parameters. As expected, the levels of normalization required varies between time points and correction is greatest for the largest magnitude peaks.

**Figure S3:**
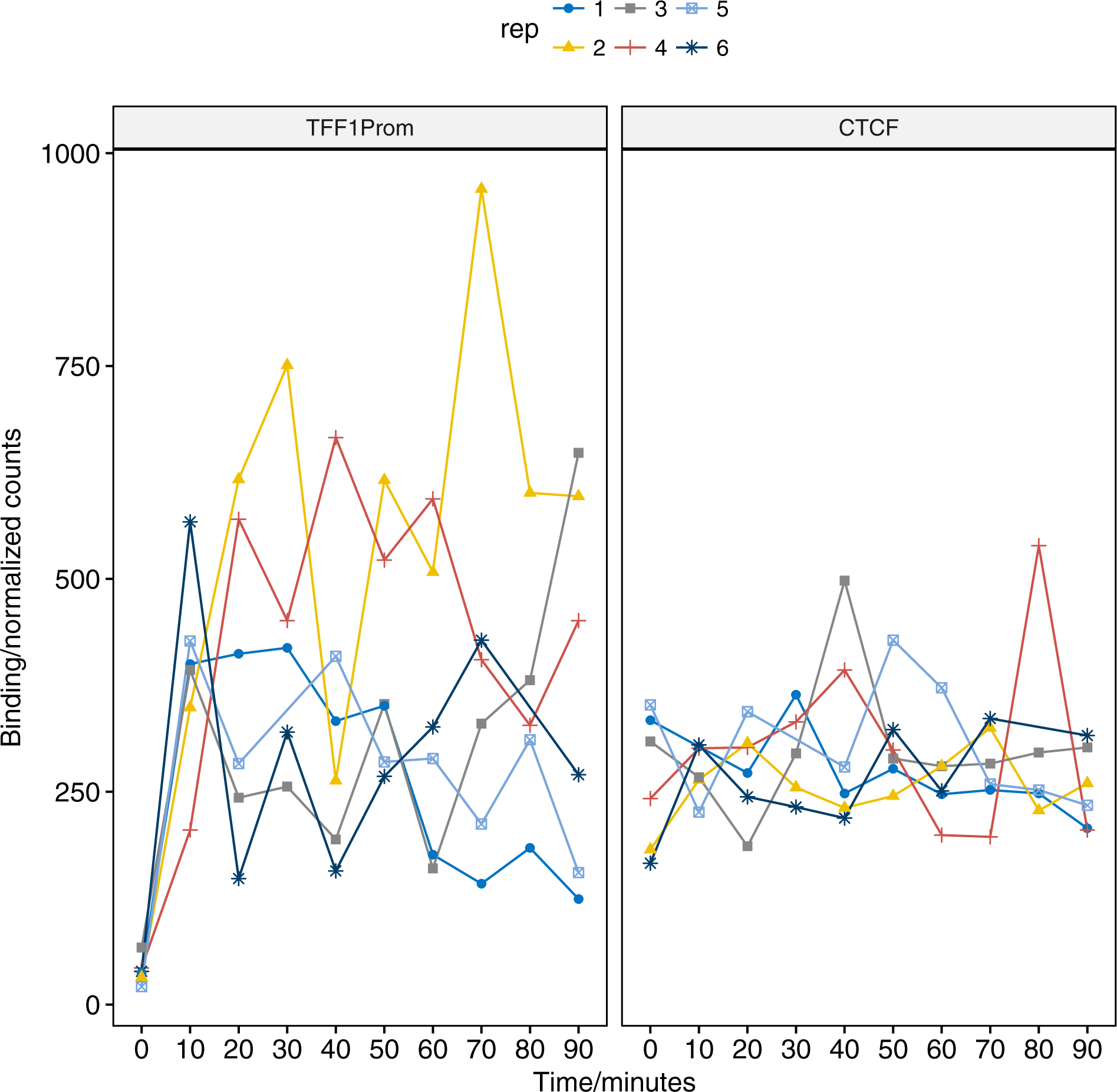
Line plot of pfChIP-seq signal at the TFF1 promoter and proximal CTCF binding site.

**Figure S4:**
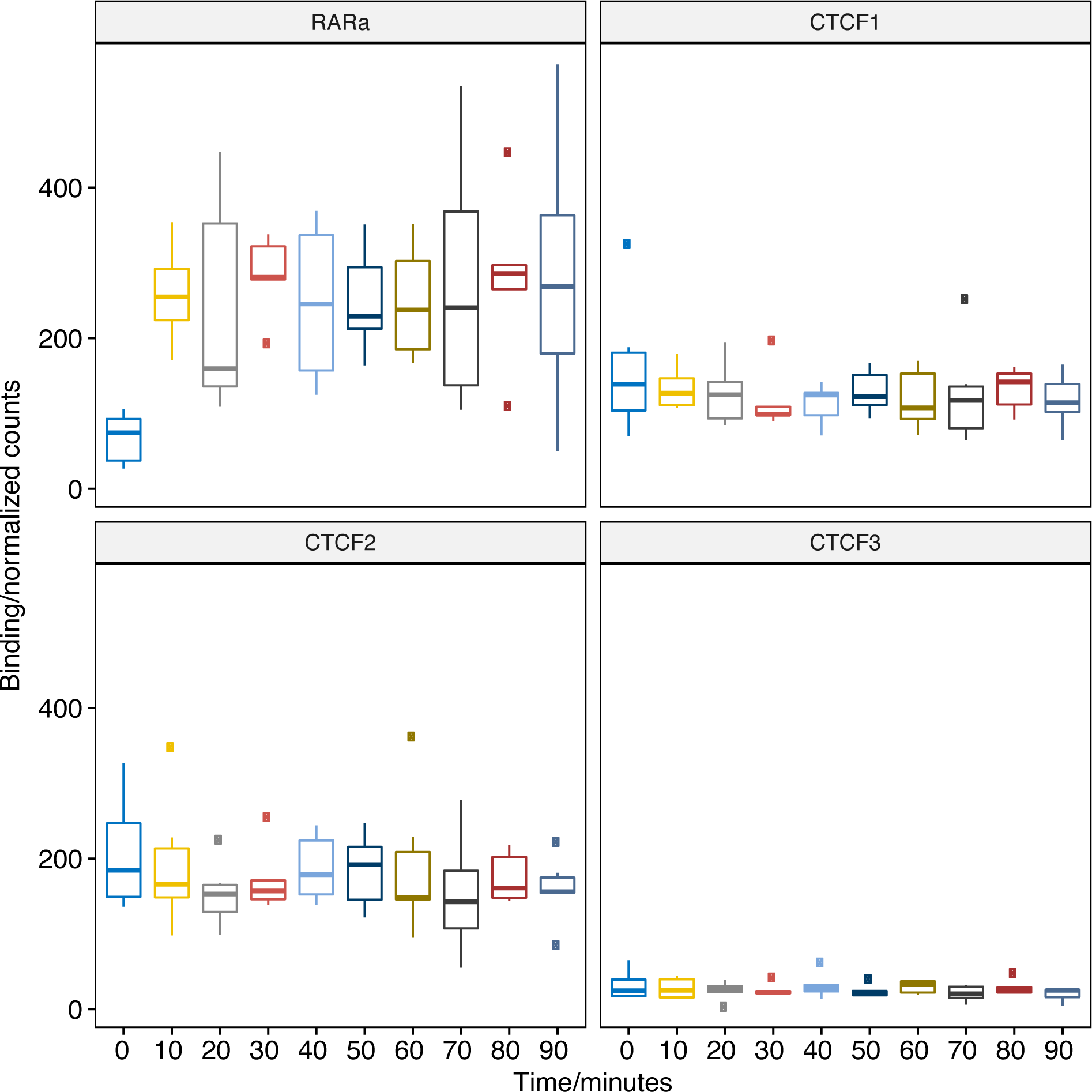
Plot of pfChIP-seq signal at the RARA promoter and proximal CTCF binding site.

**Figure S5:**
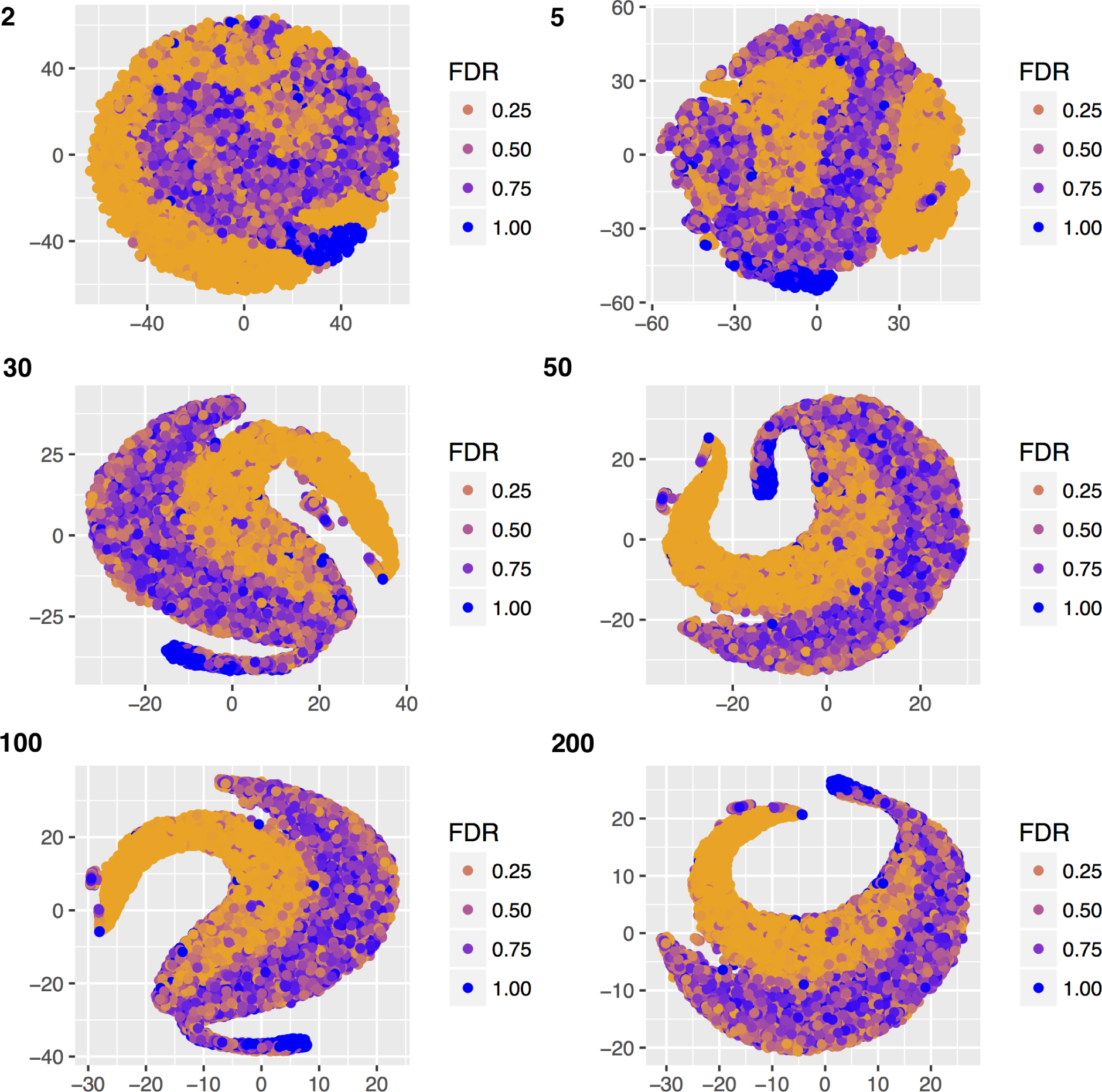
Multiple t-SNE plots of ER binding affinity changes in response to estra-2-diol at increasing perplexity (top left of each figure). For perplexity 30–200, two consistent trajectories are seen, with exact pattern depending on random seed provided. A perplexity of 200 was used to render Figure 3.

**Figure S6:**
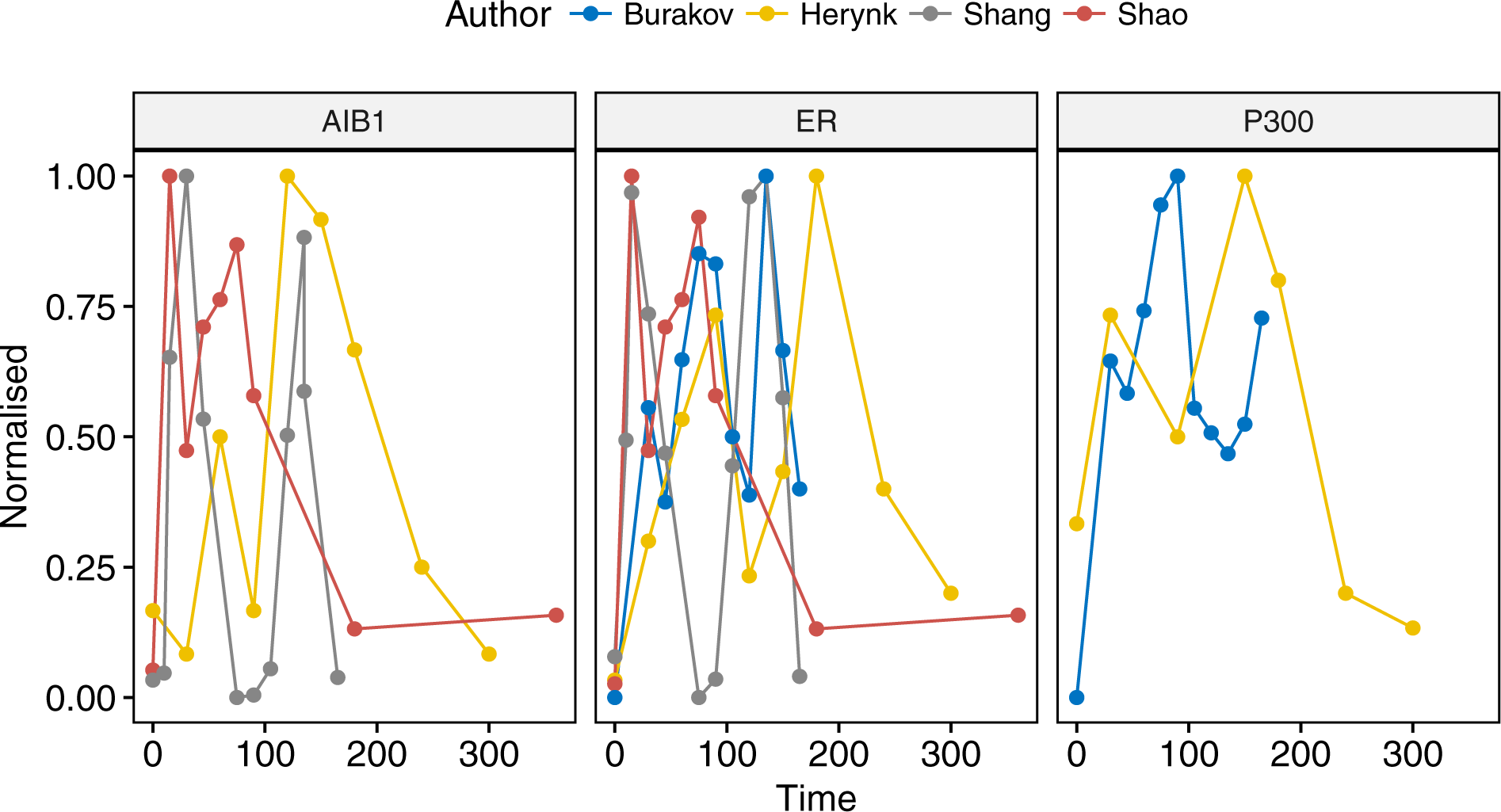
Analysis of ER and cofactor binding at the TFF1 promoter in four studies [7, 4, 1, 6]. Data was either read directly from plots within the original publication or using ImageJ [34] to calculate band density. To ensure data was comparable, data was normalized to the maximum value and all studies were chosen for replicating the conditions in Shang *et al.*’s original study.

